# The DJ-1-derived peptide, ND-13, confers cardioprotection by inhibiting mitochondrial fission and preserving mitochondrial bioenergetics via the RhoA-ROCK1-Drp1 pathway

**DOI:** 10.1101/2024.06.24.600111

**Authors:** Aishwarya Prakash, Gustavo E Crespo-Avilan, Sze Jie Loo, En Ping Yap, Jasper Chua, Ching Jianhong, Zeng Yitong, Ying-Hsi Lin, Whendy Contreras, Fan Yu, Shuo Cong, Linh Chi Dam, Chrishan J Ramachandra, Siavash Beikoghli Kalkhoran, Shengjie Lu, Daniel Offen, Sauri Hernandez-Resendiz, Derek J Hausenloy

**Affiliations:** Duke-NUS Medical School, Cardiovascular and Metabolic Disorders Programme, Singapore, Singapore; National Heart Centre Singapore, National Heart Research Institute Singapore, Singapore, Singapore; Institute of Physiological Chemistry, Technische Universität Dresden, Dresden, Germany; National University of Singapore, Yong Loo Lin School of Medicine, Singapore, Singapore; Institute of Cardiovascular Science, University College, The Hatter Cardiovascular Institute, London, UK; The Felsenstein Medical Research Center, School of Medicine, Tel Aviv University, Israel; University College London, The Hatter Cardiovascular Institute, London, UK

**Keywords:** Myocardial ischemia-reperfusion injury, mitochondrial fission, Drp1, ND-13, cardioprotection

## Abstract

**Background:** Acute myocardial infarction (AMI) and post-infarct heart failure (HF) are among the leading causes of death and disability worldwide. As such, new treatments are urgently needed to protect the heart against the detrimental effects of acute ischemia/reperfusion injury (IRI), in order to prevent the onset of HF and improve clinical outcomes following AMI. Given that mitochondrial dysfunction is a key determinant of IRI-induced cardiomyocyte death in AMI, we investigated ND-13, a 13-amino acid peptide derived from the pro-survival protein DJ-1, as a novel mitoprotective strategy for limiting myocardial infarct size (IS) following AMI.

**Methods and Results:** In isolated adult Dendra-2 mice cardiomyocytes subjected to simulated IRI, treatment with ND-13 peptide reduced cell death by 42%, decreased phosphorylation of Drp1 at Ser616 and inhibited mitochondrial fission, improved mitochondrial respiratory function, decreased oxidative stress, and preserved ATP levels. Treatment of cardiomyocytes with ND-13 decreased levels of RhoA and reduced activity of downstream ROCK1, the latter of which is known to phosphorylate Drp1 at Ser616. In *ex vivo* Langendorff-perfused hearts subjected to IRI, treatment with ND-13 at reperfusion reduced IS by 43%, attenuated oxidative stress and preserved post-infarct cardiac contractile function. Finally, *in vivo* administration of ND-13 peptide at reperfusion in mice subjected to acute myocardial IRI reduced IS by 35% at 72-hours of reperfusion, and restored mitochondrial bioenergetics in cardiomyocytes isolated from the area-at-risk following 2-hours reperfusion as evidenced by preservation of key metabolites involved in glucose oxidation and fatty acid oxidation. The cardioprotective phenotype translated to improved cardiac function and less adverse left ventricular remodelling at 28 days post-infarction.

**Conclusions:** We show for the first time the mitoprotective effects of the DJ-1-derived peptide, ND-13, administered at the onset of reperfusion following AMI. We found that ND-13 protects mitochondria by inhibiting IRI-induced mitochondrial fission and preserved mitochondrial function following IRI via the RhoA-ROCK-Drp1 pathway. These findings highlight ND-13 as a novel mitoprotective agent which has the therapeutic potential for limiting IS and preventing HF in patients with AMI.

**Clinical Perspective:** *What Is New?:*

- Administration of the DJ-1-derived peptide, ND-13, at the onset of reperfusion reduced myocardial infarct size and preserved cardiac contractile function following acute myocardial ischemia/reperfusion injury (IRI).
- The cardioprotective effect of ND-13 treatment was mediated by inhibition of IRI-induced mitochondrial fission and improvement in mitochondrial bioenergetics via the RhoA-ROCK-Drp1 pathway.

*What are the Clinical Implications?:*

- Treatment with ND-13 peptide at the onset of reperfusion has the therapeutic potential to reduce myocardial infarct size and prevent the onset of heart failure in ST-segment elevation myocardial infarction patients undergoing primary percutaneous coronary intervention.

## 1. Introduction

Acute myocardial infarction (AMI) and the heart failure (HF) that frequently follows are among the leading causes of death and disability worldwide. As such, new treatments are urgently needed to protect the heart against the detrimental effects of acute myocardial ischemia/reperfusion injury (IRI) in order to reduce myocardial infarct size (IS), preserve cardiac contractile function and prevent the onset of HF following AMI^1^. Given that mitochondrial dysfunction is a key determinant of IRI-induced cardiomyocyte death, innovative therapeutic strategies capable of preserving mitochondrial function following IRI have the potential to improve clinical outcomes following AMI^1–3^.

In this regard, changes in mitochondrial shape have been reported to impact on mitochondrial function and cell survival following IRI with elevations in intracellular calcium (Ca^2+^) and oxidative stress driving mitochondrial fission via dynamin-related protein 1 (Drp1) resulting in fragmented dysfunctional mitochondria and cardiomyocyte death during IRI^4–6^. Genetic and pharmacological ablation of Drp1 has been reported to inhibit IRI-induced mitochondrial fission and reduce IS in pre-clinical studies positioning Drp1 as a viable target for cardioprotection. Upregulation of Rho-associated coiled coil-containing protein kinase (ROCK1) has also been linked to cardiomyocyte death in preclinical models of IRI^7–9^ and inhibiting ROCK activation during IRI has been reported to exert cardioprotection in pre-clinical studies^8^. Furthermore, circulating leucocyte ROCK1 levels have been shown to be elevated in ST-segment elevation myocardial infarction (STEMI) patients^9,10^. Interestingly, recent studies have found that activation of ROCK1 induces mitochondrial fission by phosphorylating Drp1 at Ser616^7,11–14^, highlighting the role of the RhoA-ROCK1-Drp1 pathway as a target for cardioprotection.

The multifunctional protein, DJ-1, has emerged as a potential therapeutic target for both neuroprotection and cardioprotection via its known mitoprotective effects^15–18^. We have developed ND-13, a 13-amino acid peptide derived from the most conserved part of DJ-1 protein^19^, the design of which was based on its ability to confer neuroprotection against oxidative and toxic insults, both *in vitro*^20^ and *in vivo*^19,21^. Interestingly, ND-13 has been demonstrated to reduce protein levels of Rho GTPase (the upstream effector of ROCK) in focal brain ischemic injury and increase complex II activity of the electron transport chain (ETC) in neurons^21^. Although the role of ND-13 has been investigated in several models of neurodegeneration, its potential as a novel cardioprotective therapy has not yet been evaluated. In this study, we demonstrated ND-13 to be a novel mitoprotective therapy for limiting IS following acute myocardial IRI and found that ND-13 mediates its cardioprotective effect by inhibiting IRI-induced mitochondrial fission and preserving mitochondrial bioenergetics by via the RhoA-ROCK1-Drp1 pathway.

## 2. Methods

Complete data availability and comprehensive tables of the major resources can be found in the Supplementary Materials. The original datasets, detailed methodologies, and materials used in this study are available from the corresponding author upon reasonable request.

### 2.1 ND-13

ND-13, a 20-amino acid peptide derived from the 12th to 24th amino acid sequence of DJ-1 protein and attached to a cell-penetrating TAT peptide (sequence: YGRKKRR-KGAEEMETVIPVD), was produced and purified by GenScript Biotech (USA). For all *in vivo* and *ex vivo* experiments, the ND-13 lyophilized powder was reconstituted in saline and for all *in vitro* experiments, reconstituted in cell culture media. Saline (for *in vivo* and *ex vivo* experiments) or cell culture media (for *in vitro* experiments) served as the respective vehicle controls in all downstream experiments.

### 2.2 Animals

Male C57BL/6J mice and Dendra2 transgenic mice (Jackson Lab #018385) with the same genetic background as C57BL/6J mice (aged 8–14 weeks, body weight ∼24-30g) were used in compliance with ethical guidelines approved by the SingHealth IACUC (#2020/SHS/1563) for *in vitro, ex vivo*, and *in vivo* studies. All mice were housed in pathogen-free environments and maintained under non-fasting conditions to preserve normal physiological states.

### 2.3 Experimental protocol

We utilized *in vitro*, *ex vivo*, and *in vivo* models of myocardial ischemia-reperfusion injury (IRI) in this study. The detailed methodology and groups used for each IRI model are detailed in Supplementary Materials.

### 2.4 Statistical analysis

Data analysis was performed using GraphPad Prism version 10. 2. 1. Normality was assessed using the Shapiro-Wilk test to validate Gaussian distribution. Two-group comparisons were made using the unpaired, 2-tailed Student’s t-test (for data with Gaussian distribution) or Mann Whitney test (for data with non-Gaussian distribution). One-way or two-way analysis of variance (ANOVA) with Bonferroni corrections were employed for multiple group comparisons. Chi squared test was employed for contingency data. Results are expressed as mean ± SEM, with P<0.05, denoting significance.

## 3. Results

### 3.1. ND-13 treatment reduced cell death in adult mice cardiomyocytes subjected to SIRI

We first evaluated the cardioprotective effects of ND-13 peptide treatment in adult Dendra 2 (D2) mice ventricular cardiomyocytes subjected to simulated IRI (**Fig. 1A**). In a ND-13 dose-response study, we demonstrated that a dose of 12.5µM conferred significant cardioprotection following SIRI (**Supp. Fig. 1A**), reducing cell death from 64±4% in control, to 37±6% with ND-13 (p<0.01; **Figs.1B and 1C**). Incubation of adult C57BL/6J (C57) ventricular cardiomyocytes with FITC-tagged ND-13 demonstrated uptake of ND-13 within 15 minutes (**Supp. Fig. 1B**), and treatment of cardiomyocytes with ND-13 conferred no cytotoxicity for up to 24-hours incubation (**Supp. Fig. 1C**).

**Figure 1.**
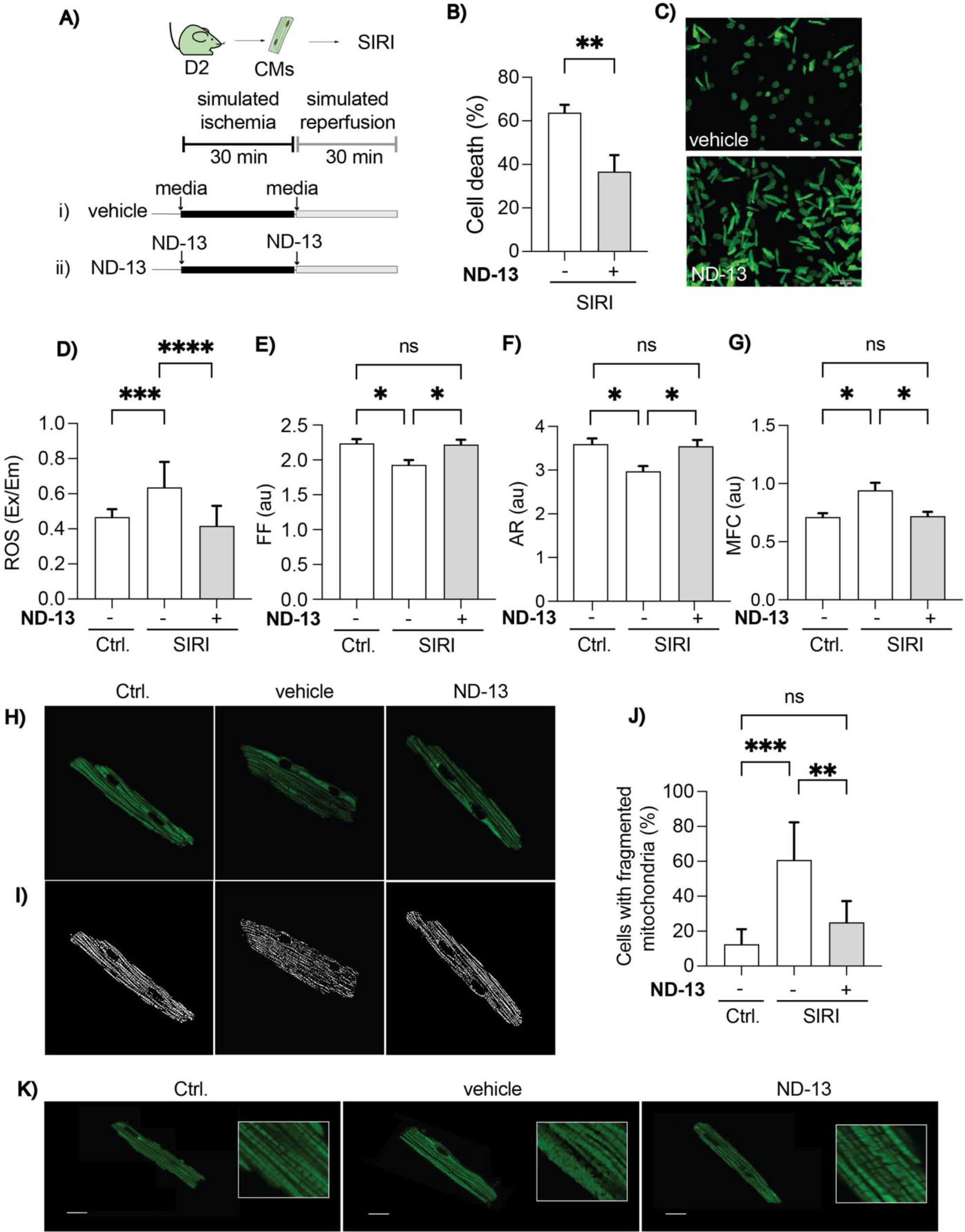
Effects of ND-13 treatment in adult Dendra2 mice cardiomyocytes subjected to SIRI. **A)** Study protocol in which ventricular cardiomyocytes isolated from Dendra2 (D2) transgenic mice were subjected to simulated ischemia/reperfusion injury (SIRI) with or without ND-13 treatment. **B)** Treatment of adult D2 mice cardiomyocytes with ND-13 significantly reduced cell death compared to the control group following SIRI. **C)** Representative confocal images of adult D2 mice cardiomyocytes following SIRI depicting reduce cell death with ND-13 treatment when compared to control. **D)** Cardiomyocyte levels of reactive oxidation species (ROS) following SIRI were significantly increased compared to control and SIRI-induced production of ROS was attenuated with ND-13 treatment. **E-G)** Mitochondrial fission assessed by was increased in adult D2 cardiomyocytes following SIRI as evidenced by decreased form factor (FF), lower aspect ratio (AT), and increased mitochondrial fragmentation count (MFC). ND-13 treatment abrogated these changes in mitochondrial morphology demonstrating that ND-13 inhibited SIRI-induced mitochondrial fission. **H-I)** Representative confocal images (fluorescence and skeletonized) of adult D2 cardiomyocytes (Vehicle) showing SIRI induced mitochondrial fission as evidenced by the presence of fragmented mitochondria when compared to cells not subjected to SIRI (Ctrl), whereas ND-13 treatment inhibited SIRI-induced mitochondrial fission and preserved the normal mitochondrial network. **J)** Quantitative analysis of the confocal images demonstrates SIRI-induced mitochondrial fission which is attenuated with ND-13 treatment. **K)** Representative zoomed-in confocal microscopic images of adult D2 cardiomyocytes showing SIRI induced mitochondrial fission as evidenced by the presence of fragmented mitochondria when compared to cells not subjected to SIRI (Ctrl). ND-13 treatment inhibited SIRI-induced mitochondrial fission and preserved the normal mitochondrial network. Abbreviations: Simulated ischemia/reperfusion injury (SIRI), Control (Ctrl), Dendra, (D2), Form factor (FF), Aspect ratio (AR), Mitochondrial fragmentation count (MFC), Reactive oxygen species (ROS), excitation/emission (Ex/Em), Cardiomyocyte (CM). (Data are expressed as mean ± SEM, N=4-15 per group, *P<0.05, **P<0.01, ***P<0.001, ****P<0.0001, by ANOVA).

### 3.2. ND-13 treatment reduced ROS production and inhibited Drp1-mediated mitochondrial fission in adult mice cardiomyocytes subjected to SIRI

We evaluated the effects of ND-13 treatment on ROS production and mitochondrial morphology in cardiomyocytes subjected to SIRI. Adult C57 ventricular cardiomyocyte levels of ROS were increased as expected following SIRI, and this effect was attenuated in the presence of ND-13 peptide treatment (0.63±0.14 Excitation/Emission (Ex/Em) in control vs. 0.40±0.11 Ex/Em with ND-13; p<0.0001; **Fig. 1D**). In adult D2 cardiomyocytes using mitochondrial two-dimensional (2D) ImageJ software analysis, we found that SIRI induced mitochondrial fission as evidenced by a decrease in form factor (FF), aspect ratio (AR) and an increase in mitochondrial fragmentation count (MFC), and these SIRI-induced effects on mitochondrial morphology were attenuated with ND-13 treatment (**Figs. 1E-1G**). Similarly, we also observed SIRI-induced mitochondrial fragmentation in cardiomyocytes using confocal microscopy to visualize individual mitochondria shape, and this effect was again attenuated by treatment with ND-13 (**Figs. 1H-1K**).

To investigate whether the inhibitory effect of ND-13 on IRI-induced mitochondrial fission was mediated by the mitochondrial fission protein, Drp1, we evaluated the effects of ND-13 on total Drp1 levels and the phosphorylation states of Ser616 (known to drive mitochondrial Drp1 translocation and fission) and Ser637 (known to prevent mitochondrial Drp1 translocation and inhibit fission). We found that SIRI reduced the total levels of Drp1 and increased the phosphorylation of Drp1 at Ser616 and these effects were attenuated in the presence of ND-13 treatment with no changes in phosphorylation of Drp1 at Ser637 (**Fig. 2A-2D**). SIRI also increased cardiomyocyte levels of Fission factor 1 (Fis1), but ND-13 treatment did not affect total Fis1 levels, and ND-13 treatment also had no significant effects on the levels of other known mitochondrial fission and fusion proteins suggesting that the inhibitory effects of ND-13 on mitochondrial fission were mediated via Drp1 (**Supp. Fig. 2).**

**Figure 2.**
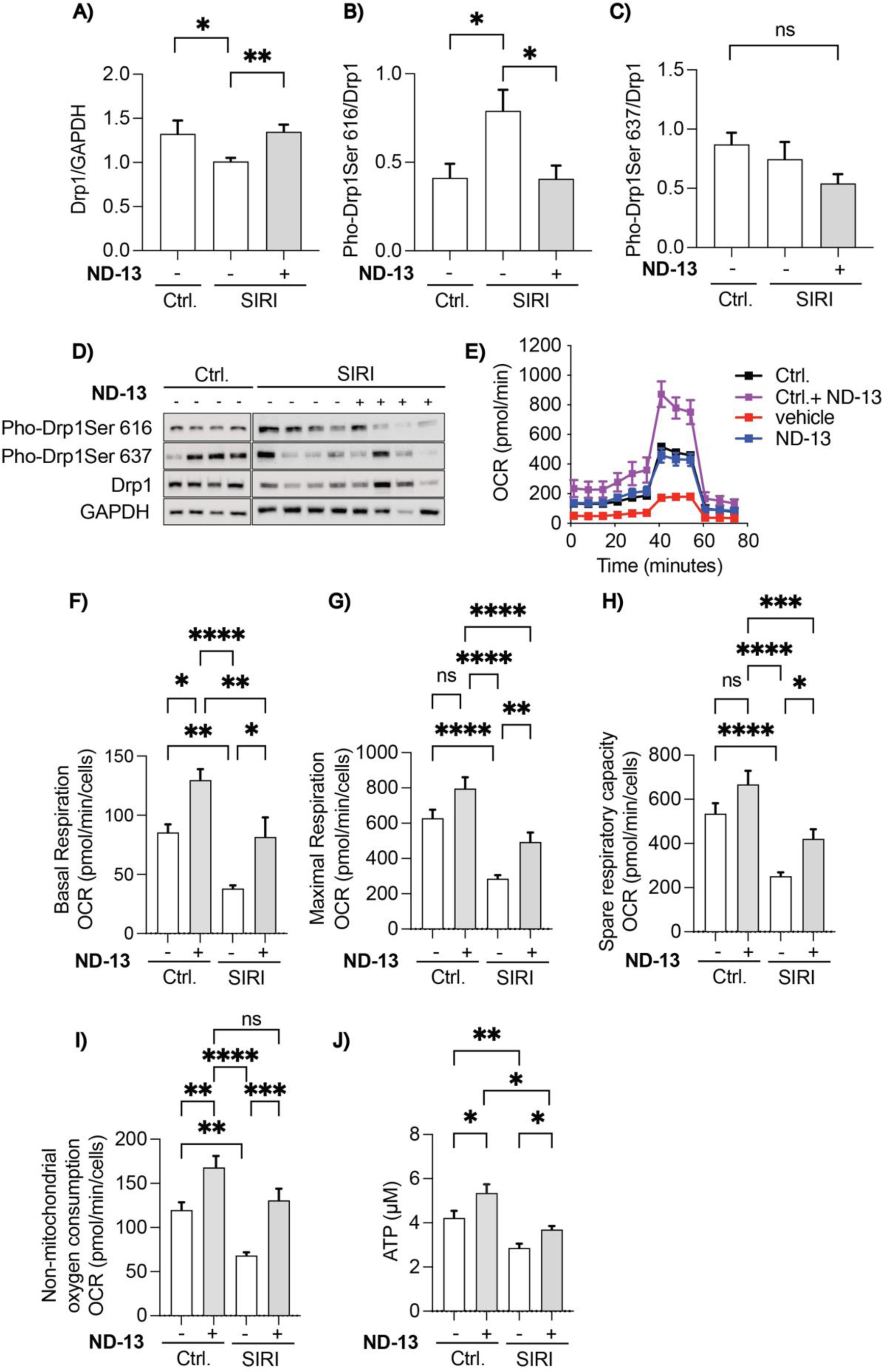
ND-13 treatment of adult D2 mice cardiomyocytes subjected to SIRI inhibits phosphorylation of Drp1 at Ser616 and preserves mitochondrial respiratory function. **A)** Total levels of Drp1 protein levels in adult D2 cardiomyocytes were significantly reduced following SIRI whereas treatment with ND-13 preserved Drp1 protein levels. **B)** Phosphorylation of Drp1 at Ser616 in adult D2 cardiomyocytes was significantly increased following SIRI whereas treatment with ND-13 a abrogated this increased. **C)** Phosphorylation of Drp1 at Ser637 in adult D2 cardiomyocytes did not alter significantly with either SIRI or ND-13 treatment. **D)** Immunoblots depicting protein levels of Pho-Drp1Ser616, Pho-Drp1Ser637 and total levels of Drp1 in adult D2 mice cardiomyocytes in control (Ctrl, no SIRI) and following SIRI in the presence and absence of ND-13 treatment. **E)** Oxygen consumption rate traces in adult C57 mice cardiomyocytes in control (no SIRI) without (Ctrl.) and with (Ctrl.+ND-13) and following SIRI in the presence (ND-13) and absence (vehicle) of ND-13 treatment. The traces show that SIRI impair mitochondrial respiration and this effect is abrogated with ND-13 treatment. **F-I)** Basal, maximal, spare respiratory capacity and non-mitochondrial oxygen consumption in adult D2 mice cardiomyocytes were all decreased following SIRI when compared to control cells (Ctrl, no SIRI), and this effect was attenuated with ND-13 treatment. **J)** ATP levels in adult D2 cardiomyocytes (vehicle) were significantly decreased following SIRI when compared to cells not subjected to SIRI (Ctrl) and treatment with ND-13 partially preserved cellular ATP levels. Abbreviations: Simulated ischemia reperfusion injury (SIRI), Control (Ctrl), Dynamin-related Protein 1 (Drp1), Serine 616 (S616), Serine 637 (S637), Phosphorylation (Pho), Glyceraldehyde 3-phosphate dehydrogenase (GAPDH), Oxygen consumption rate (OCR), Adenosine triphosphate (ATP). (Data are expressed as mean ± SEM, N=3-12,*P>0.05,**P<0.01, ***P<0.001,****P<0.0001, by ANOVA).

### 3.3. ND-13 treatment enhanced mitochondrial respiratory function in adult mice cardiomyocytes subjected to SIRI

We evaluated the effects of ND-13 treatment on mitochondrial oxygen consumption rates (OCR) in adult C57 cardiomyocytes in the presence and absence of SIRI (**Fig. 2E**). In cardiomyocytes subjected to SIRI there was a reduction in mitochondrial respiratory function as evidenced by significant reductions in: baseline and maximal respiration, spare respiratory capacity, and non-mitochondrial oxygen consumption, and these IRI-induced effects were attenuated in the presence of ND-13 demonstrating that treatment with ND-13 preserved mitochondrial respiratory function following SIRI (**Figs. 2F-2I**). Furthermore, as expected SIRI decreased levels of cardiomyocyte ATP which was restored with ND-13 treatment (**Fig. 2J**). This increased respiratory capacity induced by ND-13, was due to the true functioning of the ETC, as evidenced by increased contribution to cytosolic ATP. To ensure that increased respiration was not a result of mitochondrial uncoupling, we performed a JC-1 assay, an immediate visual readout with dual-color fluorescence (red and green), allowing the distinction between healthy (polarized) and damaged (depolarized) mitochondria. The preserved mitochondrial membrane potential after ND-13 treatment was verified, further validating the true increase in ETC function. (**Fig. 2J and Supp. Fig. 3**). Interestingly, ND-13 treated cardiomyocytes under normal conditions (non-SIRI) also experienced an increase in mitochondrial respiration, as evidenced by elevated OCR and ATP levels while maintaining an intact mitochondrial membrane potential. This underscores the capacity of ND-13 to increase mitochondrial function by improving substrate supply and the possible effect in the ETC enzymes activity (**Fig. 2E-2J, and Supp. Fig. 3**).

### 3.4. ND-13 treatment inhibited the RhoA-ROCK pathway upstream of Drp1 in adult mice cardiomyocytes subjected to SIRI

To investigate the mechanisms through which ND-13 treatment decreased the phosphorylation of Drp1-Ser616, we evaluated the effects of ND-13 on the activities of kinases which are known to regulate Drp1 at Ser616 and these included CAMKII, ERK1/2, PKCδ, GSK3β and RhoA-ROCK1 (**Supp.** Figs. 4A-4H). Western blot analysis revealed an expected trend of CAMKII and PKCδ activation in SIRI, which ND-13 treatment did not alter (**Supp. Figs. 4A-4C**). No changes in ERK1/2, GSK3β were detected at this time point with either SIRI or ND-13 treatment **(Supp. Figs. 4D-4G**). Interestingly, treatment of cardiomyocytes with ND-13 did however regulate the RhoA-ROCK1 pathway following SIRI with a significant reduction in total protein levels of RhoA with no effect on total ROCK1 levels, although the SIRI-induced increase in ROCK1 activity (assessed by phosphorylation of the ROCK1 substrate, Myosin Phosphatase subunit 1 [MYPT1] at Thr696) was significantly attenuated with ND-13 treatment (**Figs. 3A-3E**).

**Figure 3.**
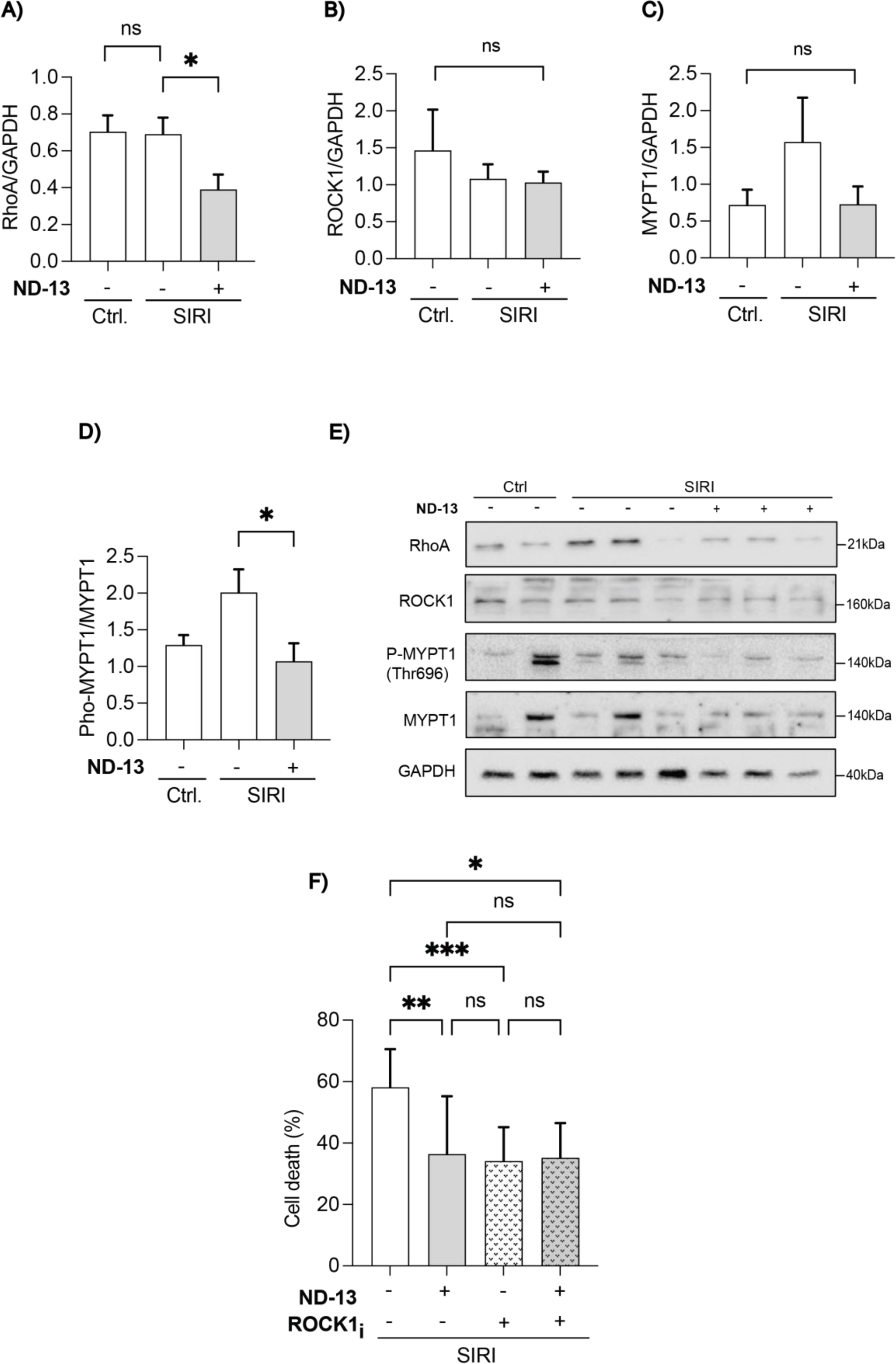
ND-13 treatment inhibited the RhoA-ROCK pathway upstream of Drp1 in adult D2 mice cardiomyocytes subjected to SIRI. **A)** Total protein levels of RhoA in adult D2 mice cardiomyocytes did not alter with SIRI but were significantly decreased with ND-13 treatment. **B)** Total protein levels of ROCK in adult D2 mice cardiomyocytes did not alter with either SIRI or ND-13 treatment. **C)** Total protein levels of MYPT1 in adult D2 mice cardiomyocytes did not alter with either SIRI or ND-13 treatment. **D)** Phosphorylation of MYPT1 at the Thr696 residue (as a marker of ROCK activity) in adult D2 mice cardiomyocytes was non-significantly increased with SIRI, and this effect was significantly decreased with ND-13 treatment. **E)** Immunoblots depicting the changes in protein levels of RhoA, ROCK, MYPT1 and phosphorylation of MYPT1-Thr696 in adult D2 mice cardiomyocytes following SIRI in the presence and absence of ND-13 treatment. **F)** Cell death in adult D2 mice cardiomyocytes following SIRI in the presence and absence of ND-13 and/or ROCK inhibitor. Treatment with either ND-13 or ROCKi alone reduced cell death following SIRI but there was no additive effect when the two treatments were combined. Abbreviations: Simulated ischemia reperfusion injury (SIRI), Control (Ctrl), Rho associated kinase 1 (ROCK1), Myosin Phosphatase Target Subunit 1 (MYPT1), Threonine 696 (Thr696), Y-27632 (ROCK1_i_), Glyceraldehyde 3-phosphate dehydrogenase (GAPDH). (Data are expressed as mean ± SEM, N=6-15 per group., *P<0.05, **P<0.01, ***P<0.001, ****P<0.0001, by ANOVA).

To further evaluate the role of ND-13 in targeting the RhoA-ROCK1 pathway, we used Y-27632, a ROCK inhibitor that attenuates phosphorylation of Drp1 at Ser616^22^ and compared the cardioprotective effects of ND-13 versus the inhibitor in cardiomyocytes subjected to SIRI.

We first undertook a dose-response study with Y-27632 and found that the dose of 5 µM conferred maximal cardioprotection and used this dose in subsequent studies (**Supp. Fig. 5**). As expected, treatment with either Y-27632 or ND-13 alone reduced cardiomyocyte death following SIRI, but interestingly when these 2 agents were administered in combination there was no additive cardioprotection suggesting that ND-13 and Y-27632 may target the same pathway i.e. RhoA-ROCK1 pathway (**Fig. 3F**).

### 3.5. ND-13 treatment reduced infarct size, preserved cardiac function, and decreased oxidative stress in *ex vivo* perfused adult mice hearts subjected to IRI

Next, we evaluated the effects of ND-13 treatment administered for the first 15 minutes of reperfusion on infarct size, contractile function and levels of oxidative stress in *ex vivo* Langendorff-perfused adult C57 mice hearts subjected to IRI (**Fig. 4A**). In a ND-13 dose-response study, we demonstrated that a dose of 20µM conferred maximal cardioprotection following IRI (**Supp. Fig. 6**), reducing IS from 53±2% in control to 29±3% with ND-13 (p<0.0001; **Figs.4B and 4C**), and preserved cardiac contractile function following IRI with improved left ventricular developed pressure (LVDP), rate pressure product (RPP) (**Figs.4D and 4E**) and preserved heart rate at constat flow rate (**Supp. Table 1**) when compared with control.

**Figure 4.**
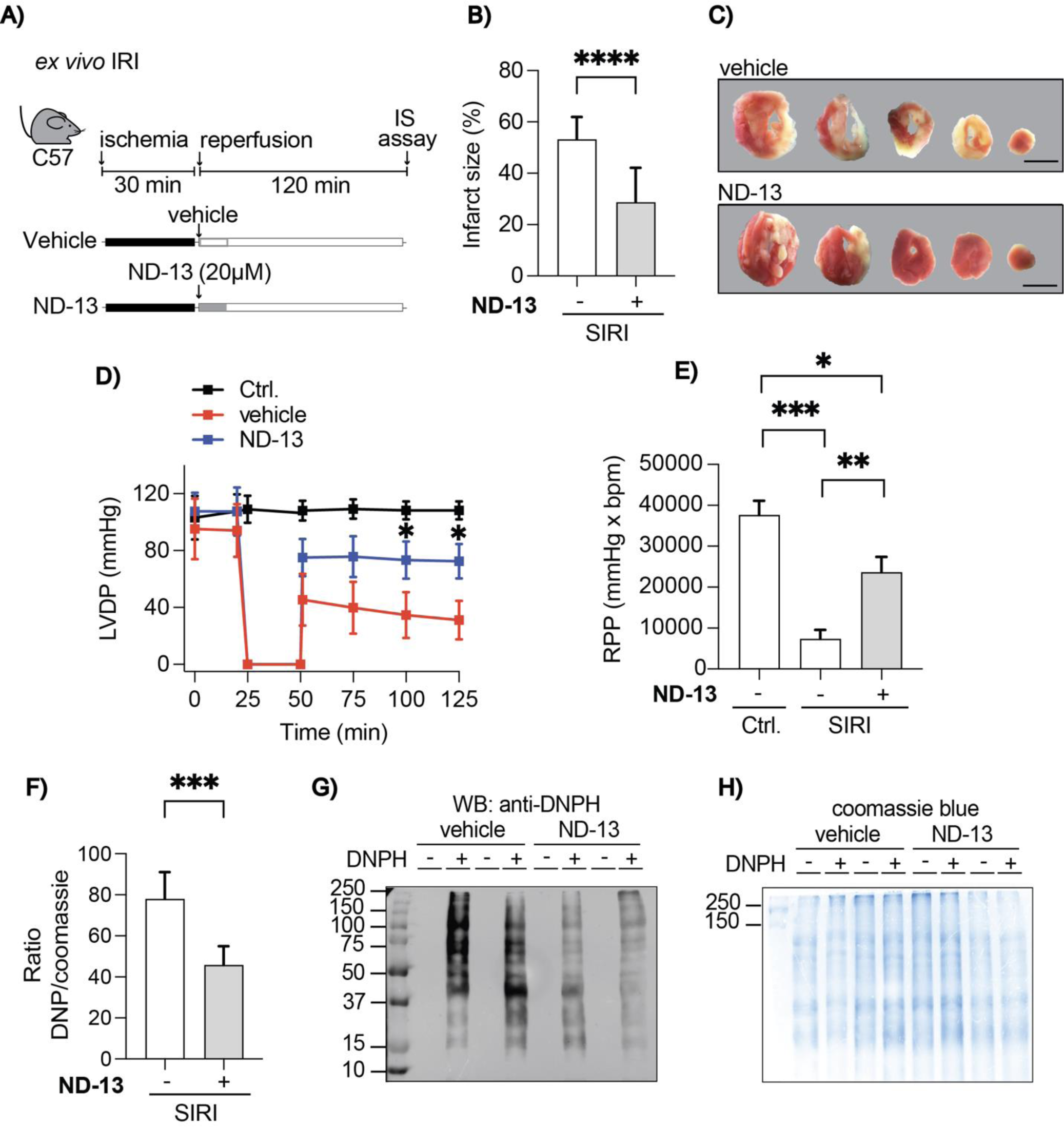
Effect of ND-13 treatment in *ex vivo* adult mice model of acute myocardial IRI. **A)** Study protocol. Hearts from C57 mice were excised and mounted on the Langendorrf-perfusion apparatus and subjected to 30 minutes of global ischemia followed by 120 minutes of reperfusion during which ND-13 or the vehicle was administered for the first 15 minutes. Infarct size (IS) was assessed by TTC staining at the end of the reperfusion period. **B)** Administration of ND-13 for the first 15 minutes of reperfusion significantly reduced infarct size following *ex vivo* IRI when compared to control. **C)** Representative heart sections with TTC staining depicting the reduced infarct size with ND-13 treatment when compared to control following *ex vivo* IRI as evidenced by a smaller unstained (pale) myocardial region compared to the vehicle control. **D)** Left ventricular developed pressure (LVDP) was measured at baseline, during global ischemia and during the reperfusion period. Compared to hearts not subjected to IRI (Ctrl), LVDP decreased with ischemia and partially recovered following ischemia. Treatment with ND-13 for the first 15 minutes of reperfusion improved the recovery of LVDP during reperfusion compared to vehicle control. **E)** Rate pressure product (RPP) calculated by the heart rate multiplied by LVDP was assessed at baseline, during global ischemia and during the reperfusion period. Compared to hearts not subjected to IRI (Ctrl), RPP decreased with ischemia and partially recovered following ischemia. Treatment with ND-13 for the first 15 minutes of reperfusion improved the recovery of RPP during reperfusion compared to vehicle control. **F)** Oxidative stress was measured using the OxyBlot protein carbonylation assay in *ex vivo* perfused C57 mice hearts following IRI. Treatment with ND-13 significantly reduced levels of DNPH-modified protein following IRI when compared to vehicle control. **G)** Representative Western blot (WB) bands with DNPH stained carbonylated proteins depicting reduced oxidative stress with ND-13 treatment. **H)** Coomassie blue staining used as a loading control for protein normalization in the WB analysis. Uniform blue staining confirmed uniform protein loaded across all groups. Abbreviations: Infarct size (IS), 2,3,5-Triphenyl-tetrazolium chloride (TTC), Left ventricular developed pressure (LVDP), Rate pressure product (RPP), Dinitrophenylhydrazine (DNPH), Area-at-risk (AAR), Infarct size (IS), Evans blue/TTC staining (EB/TTC). (Data are expressed as mean ± SEM, N=3-23 per group,*P<0.5, **P<0.01,***P<0.001, ****P<0.0001, by Students t-test).

The excessive production of ROS and the insufficiency of the antioxidant defence mechanisms leads to lipid and protein oxidation, causing cellular dysfunction and death, which is a key process driving cell death in the IRI heart ^23^. Treatment with ND-13 significantly reduced the levels of protein carbonyls, indicating a mitigation of oxidative stress in *ex vivo* perfused adult mouse hearts (**Figs. 4F-4H**).

### 3.6. ND-13 treatment reduced infarct size and preserved mitochondrial energetics in adult mice hearts subjected to *in vivo* IRI

Next, we evaluated the effects of ND-13 treatment administered 5 minutes prior to reperfusion on IS (**Fig. 5A**) and mitochondrial bioenergetics in cardiomyocytes isolated from the area-at-risk in adult C57 and D2 mice hearts subjected to *in vivo* IRI respectively. In a ND-13 dose-response study, we demonstrated that a dose of 60mg/kg conferred maximal cardioprotection following IRI (**Supp. Fig. 7**), reducing IS/AAR from 51±2% in control to 33±3% with ND-13 (p<0.0001); (**Figs. 5B-C**) with no difference in AAR between the treatment groups (**Supp. Fig. 7**). To confirm the *in vivo* myocardial uptake of ND-13 administered by intravenous injection 5 minutes before reperfusion, we demonstrated uptake of FITC-tagged ND-13 peptide into the AAR at 10 minutes of reperfusion (**Fig. 5D**).

**Figure 5.**
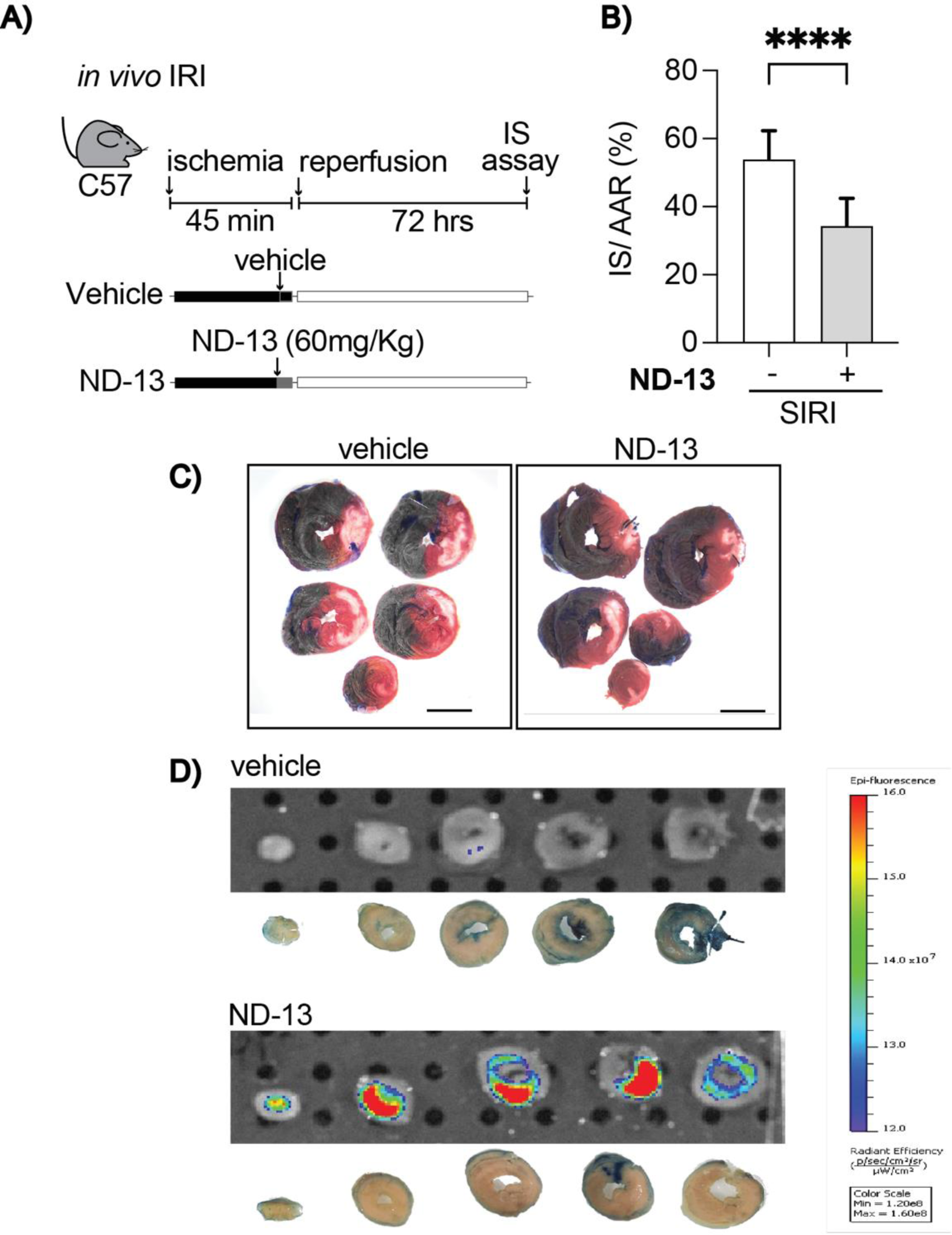
Effect of ND-13 treatment in *in vivo* adult mice model of acute myocardial IRI. **A)** Study protocol. Adult D2 mice were subjected to *in vivo* IRI with vehicle or ND-13 administered 5 min before reperfusion and infarct size/area-at-risk was determined after 72 hours using dual straining with Evans Blue and TTC. **B)** Administration of ND-13 5 minutes before reperfusion reduce myocardial infarct size (measured by IS/AAR%) when compared to vehicle control in adult C57 mice subjected to *in vivo* IRI. **C)** Representative images of EB/TTC stained heart sections indicating a visible reduction in the IS with ND-13 treatment in the *in vivo* model of IRI. EB identifies the AAR while the TTC stains the viable tissue leaving the infarcted tissue unstained and pale in color. **D)** IRI sections of the heart with saline injection observed under IVIS fluorescence imaging system (top panel) and the same sections observed under normal microscopy (bottom panel). No visible fluorescence was observed validating lack of background signal. Evans blue dye was injected to delineate the AAR. Hearts were imaging at 10 minutes of reperfusion. **E)** FITC-ND-13 treated IRI heart sections observed under the IVIS fluorescence imaging system (top panel) and the same sections under normal microscopy (bottom panel). A strong fluorescence signal was observed in the AAR of the myocardium as visualized as the area lacking the Evans blue dye. Hearts were imaged at 10 minutes reperfusion indicating the fast entry time of the peptide into the myocardium by intravenous administration. Abbreviations: Area-at-risk (AAR), Infarct size (IS), Evans blue/TTC staining (EB/TTC). (Data are expressed as mean ± SEM, N=20 per group, ****P<0.0001, by Students t-test).

We evaluated the *in vivo* effects of ND-13 treatment on the acylcarnitine profile and intermediates of the TCA cycle in cardiomyocytes harvested from within the AAR at 120 minutes reperfusion (**Figs. 6A-6F**). There was a significant and marked increase in levels of acetylcarnitine (C2) (suggesting inhibition of glucose oxidation), and a drastic decrease in certain long and very long chain acyl carnitines (Palmitoyl carnitine (C16), Stearoyl carnitine (C18), Eicasanoyl carnitine (C20), Behenoyl carnitine (C22) (suggesting inhibition of fatty acid oxidation), when compared to control hearts (**Figs. 6A-6B**). These results suggest impaired bioenergetics in cardiomyocytes from the area-at-risk region following IRI with elevated levels of acetylcarnitine that has the potential to disrupt the acetyl-CoA: acetylcarnitine ratio which in turn impacts pyruvate dehydrogenase (PDH) flux, a critical enzyme in the aerobic respiration of glucose^24^. ND-13 treatment was found to lower the levels of C2 back to that of healthy cardiomyocytes which suggests improved metabolic substrate utilization to generate energy. Long chain acyl carnitines are formed from long chain fatty acids to facilitate their transport into the mitochondria for β-oxidation, and therefore its dysregulation may indicate impaired bioenergetics^25^. Corroborating our findings to suggest impaired β-oxidation by the reduction of long chain acyl carnitines, the reduction in levels of C16 and C18 acyl carnitines have also been identified as biomarkers for impaired β-oxidation in Parkinson’s disease which further validates its’ role in fatty acid oxidation^26^. The observed increase in levels of acetyl carnitines have also been reported to inhibit the activity of Carnitine palmitoyl transferase 1 (CPT1) which catalyses the transport of long chain fatty acids into the mitochondria for β-oxidation, further validating the reduced levels of long chain acyl carnitines as a marker for impaired fatty acid oxidation^24^. The restoration of these long chain acyl carnitines to control levels with ND-13 treatment suggests an improvement in β-oxidation to fuel energy production via the TCA cycle. ND-13 treatment had no significant effect on other acyl carnitine species (**Supp. Fig. 9**).

**Figure. 6.**
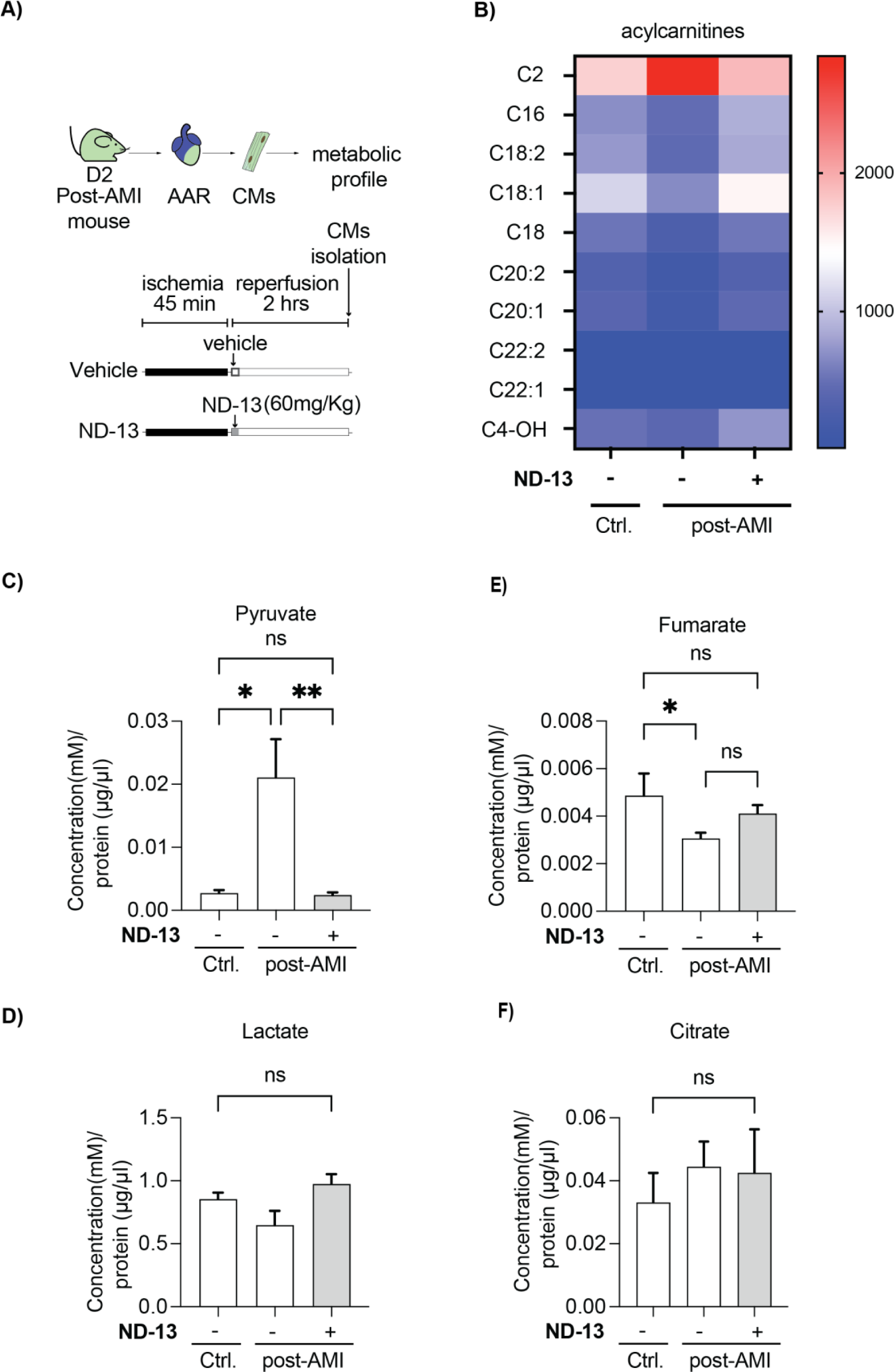
Metabolic profiles of adult ventricular cardiomyocytes harvested from the area-at-risk following *in vivo* IRI. **A)** Study design for *in vivo* IRI with and without ND-13 treatment. Dendra2 mice were subjected to IRI with one group receiving ND-13 the other with the vehicle control, 5 minutes before reperfusion. CMs from the AAR were isolated at 2 hours of reperfusion and assessed for the acyl carnitine and organic acid metabolomic profiles. **B)** The heat map depicts the significant changes in different acyl carnitine species in CMs harvested from control mice heart, and the area-at-risk of vehicle treated and ND-13 treated hearts. There was a significant accumulation of C2 acyl carnitine and reduction of the long chain and very long chain acyl carnitines in IRI. These levels were restored with ND-13 treatment indicating more efficient fatty acid oxidation to potentially produce more ATP to preserve CM viability. **C)** Levels of pyruvate are increased with IRI and attenuated to control levels with ND-13 treatment suggesting improved glucose oxidation. **D)** Lactate levels are unchanged following IRI with or without ND-13 treatment compared to the healthy control cardiomyocytes. **E)** Fumarate levels are significantly reduced upon IRI with no change with ND-13 treatment. **F)** Citrate levels are unchanged in IRI with and without ND-13 treatment compared to the control. Abbreviations: Ischemia reperfusion injury (IRI), Cardiomyocyte (CM), Area-at-risk (AAR), Adenosine triphosphate (ATP). (Data are expressed as mean ± SEM, N=3 per group, *P>0.05, **P<0.01, by ANOVA)

Following IRI, there was a significant increase in cardiomyocyte pyruvate levels which may have been due to inhibition of PDH activity from the accumulated acetylcarnitines^24^ (**Fig. 6C**). Importantly, treatment with ND-13 lowered levels of both pyruvate and acetyl carnitine, suggesting an improvement in glucose oxidation and activation of the TCA cycle. Lactate levels were unchanged across all groups suggesting no modulation of anaerobic respiration at 2 hours of reperfusion (**Fig. 6D**). Interestingly, there was a striking reduction in the cardiomyocyte levels of succinate following IRI which was preserved with ND-13 treatment although statistically analysis could not be performed as levels of succinate could not be detected across all samples in IRI (**Supp. Fig.9)**. Succinate accumulates in ischemia and its rapid oxidation following reperfusion induces reverse electron transport resulting in the release of ROS and increasing oxidative stress^27^. The preservation of succinate levels with ND-13 treatment following IRI suggests an efficient functioning of the TCA cycle which prevents the release of ROS and oxidative stress. A similar therapeutic approach using succinate dehydrogenase inhibitors to prevent the oxidation of succinate to fumarate to induce cardioprotection have also been reported which further validates this cardioprotective mechanism^28^. We also demonstrated a reduction in the levels of fumarate in cardiomyocytes following IRI suggestive of impaired oxidation of fumarate which is associated with increased cellular death, although the alleviating effects of ND-13 on fumarate levels was not significant (**Fig. 6E**). Assessing the levels of citrate, despite no changes across the groups, further provides valuable insights into the TCA cycle function (**Fig. 6F**). The levels of pyruvate and C2 acyl carnitines have been found to be elevated with IRI compared to the control, while the levels of citrate are comparable. This implies impaired TCA cycle function due to the build-up of these substrates. ND-13 treated CMs, however display lowered levels of pyruvate and C2 acyl carnitine accumulation with comparatively high levels of citrate, suggesting the forward reaction and improved function.

Finally, we evaluated the effects of ND-13 treatment administered 5 minutes prior to reperfusion in adult mice hearts subjected to *in vivo* IRI on the activation of the reperfusion injury salvage kinase (RISK) and survivor activating factor enhancement (SAFE) pathways, which are known cytoprotective signalling cascades mediating cardioprotection in the first 5 to 10 minutes of reperfusion. We found that ND-13 treatment did not activate the PI3K/AKT and MEK/ERK1/2 components of the RISK pathway or the JAK/STAT components of the SAFE pathway at 5 or 10-minutes reperfusion when compared to control (**Supp. Fig.8)**. This data provides further support for the selective role of ND-13 in regulating mitochondrial dynamics via the RhoA-ROCK1-Drp1 pathway to preserve cardiomyocyte viability.

### 3.7. Effect of ND-13 on cardiac function and fibrosis in 28 day-post-AMI hearts

A single bolus of ND-13 (60mg/kg) was found to significantly reduce IS and confer acute cardioprotection. We aimed to further assess the extent of cardioprotection conferred by this single bolus intervention in terms of preventing adverse left ventricular remodeling and preserving cardiac function across the 28 days reperfusion in a mouse model of AMI. Serial echocardiographic measurements across 1-, 7-, 28-days reperfusion revealed a significant preservation of cardiac contractile function in terms of global longitudinal strain (GLS) analysis (**Fig.7A and Supp. Table 2)**) and a trend for improvement with the less sensitive measure of cardiac function, ejection fraction (EF) (**Fig. 7B and Supp. Table 2**). However, the fibrotic scar region assessed at the end point of 28 days reperfusion, resulted in the same percentage of fibrotic scar in the tissue across the ND-13 and vehicle experimental groups (**Figs. 7C and 7D**), suggestive of positive fibrotic remodeling to preserve cardiac function. In this regard, the percentage of hearts demonstrating adverse left ventricular (LV) wall thinning was reduced from 77% with the vehicle to 30% with ND-13 treatment (p<0.05) further supporting that ND-13 treatment was able to prevent post-infarct adverse LV remodeling to maintain cardiac contractile function (**Figs. 7E**).

**Figure 7.**
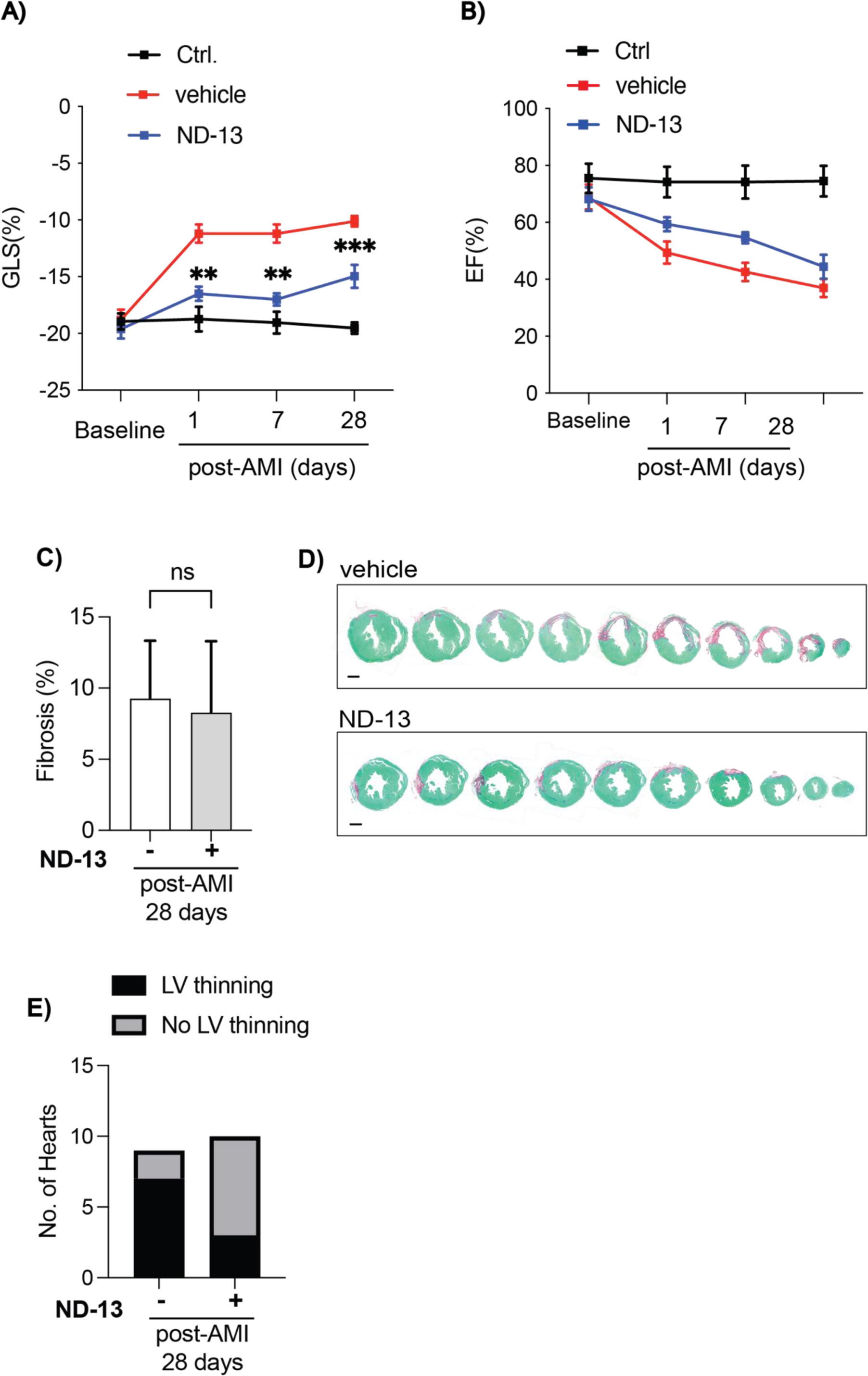
Effect of ND-13 on cardiac contractile function and myocardial fibrosis in adult mice subjected at 28 days post-infarction. **A)** Administration of ND-13 5 minutes before reperfusion in adult C57 mice subjected to *in vivo* IRI demonstrated improved cardiac contractile function as evidenced by the significant improvements in the global longitudinal strain (GLS) when compared to vehicle control. **B)** Administration of ND-13 5 minutes before reperfusion in adult C57 mice subjected to *in vivo* IRI demonstrated a trend in improvement of cardiac contractile function measured by ejection fraction (EF) when compared to vehicle control. **B)** Quantitative analysis of the fibrotic scar demonstrated no effects with ND-13 treatment compared to vehicle control. **C)** Proportion of hearts with and without LV thinning suggesting of significant post-infarct adverse LV remodelling across both groups. Only 30% ND-13-treated hearts demonstrated LV thinning compared to 77% in vehicle-treated hearts (p<0.05), suggesting that ND-13 treatment prevented post-infarct adverse LV remodelling. **D)** Representative histological heart sections at 28 days reperfusion with vehicle treatment (above panel) and ND-13 treatment (below panel) with Sirius Red/Fast Green staining. The Sirius Red stains the collagen to demarcate the fibrotic region while the fast green stains non-collagenous proteins. Abbreviations: Global longitudinal strain (GLS), Acute myocardial infarction (AMI), Left ventricle (LV), Ejection Fraction (EF). (Data are expressed as mean ± SEM, N=5-11 per group, by ANOVA in (A) Students t-test in (B) and Chi squared test for (C), scale bar represents 1mm).

## 4. Discussion

The major novel findings of our study include: (1) We show for the first time that treatment with the DJ-1-derived peptide, ND-13, reduced cell death in isolated adult mice cardiomyocytes subjected to SIRI; (2) We demonstrate that administration of ND-13 at the clinical relevant time-point of reperfusion limited myocardial IS and preserved post-infarct cardiac contractile function in adult mice hearts subjected to *ex vivo* and *in vivo* IRI; (3) We found that the cardioprotective effects of ND-13 were mediated by inhibition of IRI-induced mitochondrial fission, and preservation of mitochondrial bioenergetics via the RhoA-ROCK-Drp1-Ser616 pathway; and (4) We demonstrate preserved cardiac function by GLS and reduced LV wall thinning in ND-13 treated hearts across the 28 day remodeling period.

We designed the ND-13 peptide based on the DJ-1 protein to harness the multi-functional neuroprotective effects of the latter as a chaperone, redox-sensing, and mitoprotective agent. The role of DJ-1 in the heart has been recently investigated with studies demonstrating that DJ-1 deficient mice develop larger myocardial IS following *in vivo* IRI, a finding which was associated with increased mitochondrial fission and impaired mitochondrial function^18,29^. More recently, the administration of recombinant DJ-1 by intraperitoneal injection 60 min prior to infarction in an *in vivo* adult mice model of IRI reduced myocardial IS, leukocyte infiltration, apoptosis and oxidative stress^30^. In the clinical setting, DJ-1 protein levels have been found to be drastically reduced in the hearts of patients with end-stage heart failure further implicating DJ-1 as a potential therapeutic target for alleviating cardiac disease^31^.

In our study, we demonstrated for the first time that treatment with the DJ-1 protein-derived peptide, ND-13, protected isolated adult mice cardiomyocytes from SIRI, and its administration at the clinically relevant timepoint of reperfusion limited myocardial IS and preserved cardiac contractile function in *ex vivo* adult mice hearts subjected to IRI. Importantly, ND-13 administered 5 minutes prior to reperfusion limited myocardial IS in adult mice subjected to *in vivo* IRI, highlighting the therapeutic potential of ND-13 peptide treatment a novel cardioprotective therapy for limiting infarct size and preventing HF following AMI. Having demonstrated less cardiomyocyte death and reduced myocardial IS with ND-13 treatment following IRI, we next investigated the mechanisms underlying ND-13 cardioprotection. Interestingly, the neuroprotective effect of ND-13 against focal cerebral ischemic injury was found to be preserved in DJ-1 deficient mice suggesting a DJ-1 independent mechanism of protection^21^, suggesting that ND-13 may also confer direct cardioprotection independent of the DJ-1 protein.

The inhibition of Drp1-mediated mitochondrial fission in myocardial IRI has garnered great attention in recent years as a potential therapeutic strategy to mitigate mitochondrial dysfunction and reduce cardiomyocyte death. There are several pharmacological inhibitors of Drp1 such as Mdivi-1 ^32–34^ and a more newly described molecule DRP1i27^35^ to inhibit Drp1 activity by obstructing GTP hydrolysis and in turn preventing mitochondrial fission. However, Midivi-1 has several off-target effects which have contributed to contradictory findings and questionable mechanistic insights^9,36^. The Drp1 inhibitor, DRP1i27 shows promise to specifically bind and inhibit Drp1 mediated mitochondrial fission. This, however, was tested in HL-1 cells, a mouse derived atrial tumor cell line which has limitations in studying mitochondrial dynamics (such as lower mitochondrial content, glycolytic metabolic preference, and lowered ATP synthesis) following IRI^37^. Additionally, these studies assessed mitochondrial morphology but failed to determine if the improved morphology affected mitochondrial function. The overexpression of DJ-1 in HL-1 cardiac cells subjected to SIRI reduced cardiomyocyte death, inhibited IRI-induced opening of the mitochondrial permeability transition pore (mPTP) and prevented mitochondrial fission, highlighting a potential role of mitochondrial morphology as a target for DJ-1 mediated cardioprotection^18^.

In our study, we demonstrated for the first time that treatment with ND-13, inhibited Drp1-mediated mitochondrial fission in isolated adult mice cardiomyocytes subjected to SIRI. Using confocal microscopic imaging of individual mitochondria in Dendra2 cardiomyocytes, we were able to demonstrate that ND-13 inhibited SIRI-induced mitochondrial fission as evidenced by decreased numbers of rounded fragmented mitochondria and preservation of the mitochondrial network. The maintenance of the normal mitochondrial network with ND-13 treatment was associated with less oxidative stress and preserved mitochondrial bioenergetics post-SIRI. The mechanism underlying ND-13-mediated inhibition of SIRI-induced mitochondrial fission was linked to the decreased phosphorylation of Drp1 at Ser616, a post-translational modification of Drp1 that is known to drive mitochondrial fission. The effect of ND-13 on mitochondrial fission appeared to be mediated by Drp1 at Ser616, as no effects were observed on the phosphorylation of Drp1 at Ser637 nor were there any effects on the expression levels of the other mitochondrial fission proteins (Mff, Fis1, Mid49, Mid51) or fusion proteins (Opa1, Mfn1, Mfn2). ND-13 treatment also preserved total Drp1 levels which may have been due to it preventing the degradation of Drp1 protein dysregulated mitophagy^38^.

Phosphorylation of Drp1 at the Ser616 residue has been reported to be catalyzed by PKC-δ, cyclin-dependent kinase 1 (CDK1), ERK1/2, calmodulin-dependent protein kinase II (CaMKII), PINK1 (PTEN-induced putative kinase 1), AMPK (adenosine monophosphate-activated protein kinase), GSK3β and by the newly reported RhoA GTPase/Rho-associated protein kinase (ROCK1) pathway^39–42^. We found that treatment of cardiomyocytes subjected to SIRI with ND-13 did not affect the expression levels of upstream kinases, PKC-δ, ERK1/2, CaMKII, or GSK3β, but interestingly, ND-13 did modulate the Rho-A-ROCK pathway. Interestingly, the neuroprotective effect of ND-13 in regulating mitochondrial function and oxidative stress has been associated with reducing protein expression of Rho GTPase 2 (a member of the Rho GTPase subfamily) therefore providing the antecedent to assess its potential cardioprotective effects in regulating the RhoA-ROCK1-Drp1 pathway to confer cardioprotection. RhoA is a small GTPase that was initially reported to mediate cytoskeletal dynamics along with its downstream effector protein kinase-ROCK1/2, with more recent studies shedding light on its role in cardiomyocyte apoptosis, fibrosis, platelet activation, leukocyte activation and endothelial dysfunction ^43^. Clinical studies further demonstrated that IRI increased ROCK activity in leukocytes collected from blood samples in patients with STEMI during the ischemic episode and at the early phases of reperfusion which corresponded to worsened prognosis ^9^. The detrimental effects of ROCK activity, in particular the ROCK1 isoform was demonstrated in a recent study by Li et al., confirming that increased ROCK1 activity as a consequence of IRI, resulted in Drp1-Ser616 driven mitochondrial fission contributing to mitochondrial dysfunction resulting in cellular death 39. In our study, we demonstrated for the first time that ND-13 treatment reduced total protein levels of the small GTPase, RhoA, which in turn reduced its ability to activate its downstream kinase, ROCK1 which selectively phosphorylates Drp1 at the Ser616 residue to initiate mitochondrial fission ^44^. Despite no changes in the total protein levels of ROCK1, we observed the reduction in its activity through the reduced phosphorylation of MYPT1 at the Thr696 residue. As a well characterized substrate of ROCK1, the phosphorylation status of MYPT1 is directly proportional to the activity of this kinase ^45^. These findings supported our proposal that ND-13 attenuates the RhoA-ROCK1-Drp1-Ser616 pathway inducing mitochondrial fission in SIRI therefore preserving mitochondrial dynamics and restoring bioenergetics. To confirm and validate the detrimental role of the ROCK1 kinase, we demonstrated that inhibition of ROCK1 activity using the ROCK1 inhibitor, Y-27632, preserved cell viability following SIRI. Additionally, the combination of ND-13 and ROCK1 inhibitor did provide additive cardioprotection in cardiomyocytes subjected to SIRI suggesting that these two agents target the RhoA-ROCK pathway.

In contrast to our findings, it has been demonstrated in another study that the activation of the RhoA-ROCK pathway increased mitochondrial fission and enhanced cell viability post-SIRI ^44^. The key reason for this contradictory result is likely due to the cell type, with their study using neonatal cardiomyocytes which have altered mitochondrial and metabolic profiles in terms of poorly aligned mitochondrial cristae resulting in lowered ETC function, and therefore predominantly relying on glycolysis for ATP production compared to adult cardiomyocytes used in our study, where the primary source of ATP is mitochondrial respiration ^46^. With this preferred method of ATP production in neonatal cardiomyocytes, increased mitochondrial fragmentation may facilitate increased glycolysis to aid in preservation of cell viability following IRI ^47^. However, in a more physiological model of SIRI with adult cardiomyocytes which depend on glucose oxidation, fatty acid oxidation and mitochondrial ETC activity for ATP production, excessive mitochondrial fragmentation induced by the RhoA-ROCK1-Drp1-Ser616 pathway would prove detrimental, and the reverse would then be protective, as seen in our study. This highlights the critical link between mitochondrial morphology and metabolic substrate utilization in SIRI which underscores the importance of therapeutic strategies to target both for maximum cellular protection.

The preservation of the mitochondrial network along with the attenuation of oxidative stress in ND-13-treated cardiomyocytes subjected to SIRI, suggested improvements in mitochondrial bioenergetics ^1^. This was supported by findings showing that ND-13 treatment of cardiomyocytes subjected to SIRI preserved mitochondrial respiration, mitochondrial membrane potential and ATP levels. To evaluate the *in vivo* metabolic effects of ND-13 following IRI, for the first time we assessed the key metabolites involved in fatty acid oxidation (acyl carnitines), and the TCA cycle (organic acids) in cardiomyocytes specifically harvested from the area-at-risk region in adult mice subjected to IRI. We found evidence of IRI-induced metabolic perturbation with elevated levels of C2 acyl carnitines, reduced levels of long chain acyl carnitines, increased levels of pyruvate and lowered levels of succinate. ND-13 treatment was found to preserve the levels of all these metabolites to that of the control indicating prevention of metabolic impairment following IRI, therefore preserving cell viability and limiting IS development.

The RISK and SAFE pathways are well described pro-survival pathways the activation of which in the first few minutes of reperfusion confer cardioprotection ^48^. In our study, we also evaluated the role of these two survival cascades in ND-13 mediated cardioprotection in adult mice subjected to *in vivo* acute myocardial IRI. Although, IRI induced the phosphorylation of AKT, ERK and STAT3, treatment with ND-13 at the onset of reperfusion did not further enhance the activation of these kinase components of the RISK and SAFE pathways. These findings suggest that ND-13 confers its cardioprotective effects independent of the RISK and SAFE pathways providing further evidence for the RhoA-ROCK-Drp1-Ser616 pathway as the target for ND-13-mediated cardioprotection.

Finally, we assessed the effect of ND-13 on cardiac function and fibrosis during the 28-day remodeling period. In our assessment of cardiac function using echocardiography, we observed a significant improvement in GLS across all days of reperfusion with ND-13 treatment. However, despite the treatment not affecting the total fibrotic scar area, the fibrosis developed appeared to contribute to better cardiac remodeling as seen through reduced LV thinning therefore supporting cardiac function. Beyond assessing the spatial distribution of fibrosis, it is also crucial to assess the composition of collagens and the associated extracellular matrix (ECM) within the fibrotic scar, as they play a significant role in organ function^49^. In the setting of myocardial infarction, the ratio of collagen I, III, and V to collagen IV is much higher, resulting in the loss of structure and function of the left ventricular wall^50^. However, if this ratio of collagen deposition is altered, it may provide an improved compensatory mechanism, even without reductions in the total fibrotic scar area. Therefore, future studies will aim to assess the composition of deposited collagen to better understand the intricacies of cardiac remodelling and its association with cardiac function. Our study highlights that preserving mitochondrial homeostasis at the onset of reperfusion lays the foundation for improved cardiac function over the 28-day reperfusion period.

## 5. Limitations and future directions

Firstly, it must be acknowledged that our studies focused exclusively on the cardioprotective role of ND-13 in adult cardiomyocytes and its effect in other non-cardiac cells such as endothelial cells, immune cells and fibroblasts were not assessed. Secondly, the mechanism through which ND-13 reduced total levels of RhoA were not investigated and may be due to ND-13 binding to and activating E3 ligases that catalyze RhoA ubiquitination for degradation or by activation of the endosomal-lysosomal/autophagy-lysosomal pathway. Thirdly, we cannot exclude other mechanisms contributing the cardioprotection induced by ND-13 given that other studies have neuroprotective effects associated with Nrf2 activation and regulation of other mitochondrial proteins (such as Rho GTPase2 and Ubiquitin Specific Peptidase 35). Finally, one may consider encapsulating ND-13 in nanoparticles to protect the peptide from protease degradation thereby increasing the bioavailability and delivery of ND-13 peptide to the ischemic heart following AMI.

In summary, we have shown for the first time that the administration of the DJ-1-derived peptide, ND-13, at the onset of reperfusion reduced myocardial IS and preserved cardiac contractile function following acute myocardial IRI. The cardioprotective effect of ND-13 treatment was mediated by inhibition of IRI-induced mitochondrial fission and improvement in mitochondrial bioenergetics via the RhoA-ROCK-Drp1 pathway (**Fig. 8**). Treatment with ND-13 peptide at the onset of reperfusion has the therapeutic potential to reduce myocardial IS and prevent the onset of HF in STEMI patients undergoing PPCI.

**Figure 8.**
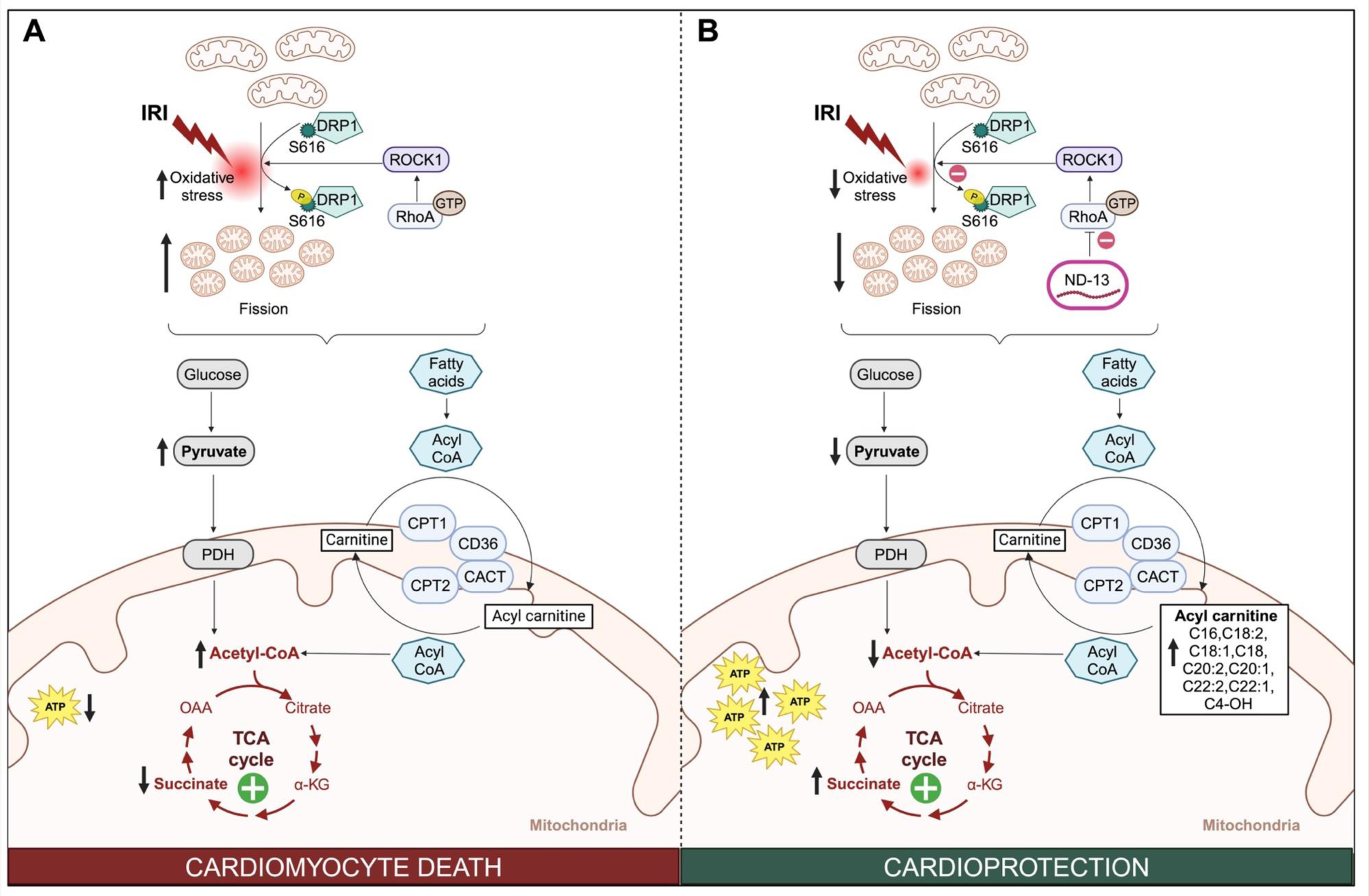
Treatment with ND-13 confers cardioprotection by inhibiting mitochondrial fission and preserving mitochondrial bioenergetics via inhibiting the RhoA-ROCK1-Drp1-Ser616 pathway. **A)** The RhoA-ROCK1-Drp1-Ser616 pathway is activated during the early reperfusion phase, plays a crucial role in inducing excessive mitochondrial fragmentation in cardiomyocytes following IRI. This leads to impaired mitochondrial function and contributes to oxidative stress. The disruption of mitochondrial homeostasis, as evidenced by the accumulation of C2 acyl carnitine and pyruvate and reduced levels of long and very long chain acyl carnitines, indicates impaired glucose and fatty acid oxidation, respectively. **B)** The therapeutic potential of ND-13 is evident in its ability to inhibit IRI-induced mitochondrial fission by targeting the RhoA-ROCK1-Drp1-Ser616 pathway. This effect preserves mitochondrial morphology and restore mitochondrial function, leading to a reduction in oxidative stress. The restoration of bioenergetics within the cardiomyocytes is demonstrated by the reduced levels of C2 acyl carnitines and pyruvate, and the elevated levels of long and very long chain acyl carnitines, suggesting restored glucose and fatty acid oxidation respectively. Abbreviations: Ischemia-reperfusion injury (IRI), Dynamin-related protein 1 (DRP1), Serine 616 (S616), RhoA-associated kinase 1 (ROCK1), Pyruvate dehydrogenase (PDH), alpha-ketoglutaric acid (a-KG), Oxaloacetic acid (OAA), Adenosine triphosphate (ATP), Carnitine palmitoyltransferase 1 (CPT1), Carnitine-acylcarnitine translocase (CACT), Carnitine palmitoyltransferase 2 (CPT2). *Note: Figure created with Biorender*.

## References

1. Hernandez-Resendiz S, Prakash A, Loo SJ, Semenzato M, Chinda K, Crespo-Avilan GE, Dam LC, Lu S, Scorrano L, Hausenloy DJ. Targeting mitochondrial shape: at the heart of cardioprotection. Basic Res Cardiol. 2023;118:49. doi: 10.1007/s00395-023-01019-9

2. Taegtmeyer H, Young ME, Lopaschuk GD, Abel ED, Brunengraber H, Darley-Usmar V, Des Rosiers C, Gerszten R, Glatz JF, Griffin JL, et al. Assessing Cardiac Metabolism: A Scientific Statement From the American Heart Association. Circ Res. 2016;118:1659–1701. doi: 10.1161/RES.0000000000000097

3. Zuurbier CJ, Bertrand L, Beauloye CR, Andreadou I, Ruiz-Meana M, Jespersen NR, Kula-Alwar D, Prag HA, Eric Botker H, Dambrova M, et al. Cardiac metabolism as a driver and therapeutic target of myocardial infarction. J Cell Mol Med. 2020;24:5937–5954. doi: 10.1111/jcmm.15180

4. Chouchani ET, Pell VR, Gaude E, Aksentijevic D, Sundier SY, Robb EL, Logan A, Nadtochiy SM, Ord ENJ, Smith AC, et al. Ischaemic accumulation of succinate controls reperfusion injury through mitochondrial ROS. Nature. 2014;515:431–435. doi: 10.1038/nature13909

5. Lucas DT, Szweda LI. Cardiac reperfusion injury: aging, lipid peroxidation, and mitochondrial dysfunction. Proc Natl Acad Sci U S A. 1998;95:510–514. doi: 10.1073/pnas.95.2.510

6. Paradies G, Petrosillo G, Pistolese M, Di Venosa N, Serena D, Ruggiero FM. Lipid peroxidation and alterations to oxidative metabolism in mitochondria isolated from rat heart subjected to ischemia and reperfusion. Free Radic Biol Med. 1999;27:42–50. doi: 10.1016/s0891-5849(99)00032-5

7. Bao W, Hu E, Tao L, Boyce R, Mirabile R, Thudium DT, Ma XL, Willette RN, Yue TL. Inhibition of Rho-kinase protects the heart against ischemia/reperfusion injury. Cardiovasc Res. 2004;61:548–558. doi: 10.1016/j.cardiores.2003.12.004

8. Patel P, Parikh M, Shah H, Gandhi T. Inhibition of RhoA/Rho kinase by ibuprofen exerts cardioprotective effect on isoproterenol induced myocardial infarction in rats. Eur J Pharmacol. 2016;791:91–98. doi: 10.1016/j.ejphar.2016.08.015

9. Zhang J, Xu F, Liu XB, Bi SJ, Lu QH. Increased Rho kinase activity in patients with heart ischemia/reperfusion. Perfusion. 2019;34:15–21. doi: 10.1177/0267659118787432

10. Li X, Wu X, Li H, Chen H, Wang Y, Li W, Ding X, Hong X. Increased Rho kinase activity predicts worse cardiovascular outcome in ST-segment elevation myocardial infarction patients. Cardiol J. 2016;23:456–464. doi: 10.5603/CJ.a2016.0031

11. Zhou Q, Liao JK. Rho kinase: an important mediator of atherosclerosis and vascular disease. Curr Pharm Des. 2009;15:3108–3115. doi: 10.2174/138161209789057986

12. Ocaranza MP, Gabrielli L, Mora I, Garcia L, McNab P, Godoy I, Braun S, Cordova S, Castro P, Novoa U, et al. Markedly increased Rho-kinase activity in circulating leukocytes in patients with chronic heart failure. Am Heart J. 2011;161:931–937. doi: 10.1016/j.ahj.2011.01.024

13. Nohria A, Grunert ME, Rikitake Y, Noma K, Prsic A, Ganz P, Liao JK, Creager MA. Rho kinase inhibition improves endothelial function in human subjects with coronary artery disease. Circ Res. 2006;99:1426–1432. doi: 10.1161/01.RES.0000251668.39526.c7

14. Calo LA, Vertolli U, Pagnin E, Ravarotto V, Davis PA, Lupia M, Naso E, Maiolino G, Naso A. Increased rho kinase activity in mononuclear cells of dialysis and stage 3-4 chronic kidney disease patients with left ventricular hypertrophy: Cardiovascular risk implications. Life Sci. 2016;148:80–85. doi: 10.1016/j.lfs.2016.02.019

15. Kaneko Y, Shojo H, Burns J, Staples M, Tajiri N, Borlongan CV. DJ-1 ameliorates ischemic cell death in vitro possibly via mitochondrial pathway. Neurobiol Dis. 2014;62:56–61. doi: 10.1016/j.nbd.2013.09.007

16. Tajiri N, Borlongan CV, Kaneko Y. Cyclosporine A Treatment Abrogates Ischemia-Induced Neuronal Cell Death by Preserving Mitochondrial Integrity through Upregulation of the Parkinson’s Disease-Associated Protein DJ-1. CNS Neurosci Ther. 2016;22:602–610. doi: 10.1111/cns.12546

17. Shimizu Y, Lambert JP, Nicholson CK, Kim JJ, Wolfson DW, Cho HC, Husain A, Naqvi N, Chin LS, Li L, et al. DJ-1 protects the heart against ischemia-reperfusion injury by regulating mitochondrial fission. J Mol Cell Cardiol. 2016;97:56–66. doi: 10.1016/j.yjmcc.2016.04.008

18. Dongworth RK, Mukherjee UA, Hall AR, Astin R, Ong SB, Yao Z, Dyson A, Szabadkai G, Davidson SM, Yellon DM, et al. DJ-1 protects against cell death following acute cardiac ischemia-reperfusion injury. Cell Death Dis. 2014;5:e1082. doi: 10.1038/cddis.2014.41

19. Lev N, Barhum Y, Ben-Zur T, Aharony I, Trifonov L, Regev N, Melamed E, Gruzman A, Offen D. A DJ-1 Based Peptide Attenuates Dopaminergic Degeneration in Mice Models of Parkinson’s Disease via Enhancing Nrf2. PLoS One. 2015;10:e0127549. doi: 10.1371/journal.pone.0127549

20. Lev N, Barhum Y, Lotan I, Steiner I, Offen D. DJ-1 knockout augments disease severity and shortens survival in a mouse model of ALS. PLoS One. 2015;10:e0117190. doi: 10.1371/journal.pone.0117190

21. Molcho L, Ben-Zur T, Barhum Y, Offen D. DJ-1 based peptide, ND-13, promote functional recovery in mouse model of focal ischemic injury. PLoS One. 2018;13:e0192954. doi: 10.1371/journal.pone.0192954

22. Shen YL, Shi YZ, Chen GG, Wang LL, Zheng MZ, Jin HF, Chen YY. TNF-alpha induces Drp1-mediated mitochondrial fragmentation during inflammatory cardiomyocyte injury. Int J Mol Med. 2018;41:2317–2327. doi: 10.3892/ijmm.2018.3385

23. Arreguin F, Garcia N, Hernandez-Resendiz S, Buelna-Chontal M, Correa F, Olin-Sandoval V, Medina-Campos ON, Pedraza-Chaverri J, Zazueta C. Attenuation of oxidant damage in the postconditioned heart involves non-enzymatic response and partial catalytic protection. Exp Physiol. 2012;97:1119–1130. doi: 10.1113/expphysiol.2012.065763

24. Schroeder MA, Atherton HJ, Dodd MS, Lee P, Cochlin LE, Radda GK, Clarke K, Tyler DJ. The cycling of acetyl-coenzyme A through acetylcarnitine buffers cardiac substrate supply: a hyperpolarized 13C magnetic resonance study. Circ Cardiovasc Imaging. 2012;5:201–209. doi: 10.1161/CIRCIMAGING.111.969451

25. McCann MR, George De la Rosa MV, Rosania GR, Stringer KA. L-Carnitine and Acylcarnitines: Mitochondrial Biomarkers for Precision Medicine. Metabolites. 2021;11. doi: 10.3390/metabo11010051

26. Saiki S, Hatano T, Fujimaki M, Ishikawa KI, Mori A, Oji Y, Okuzumi A, Fukuhara T, Koinuma T, Imamichi Y, et al. Decreased long-chain acylcarnitines from insufficient beta-oxidation as potential early diagnostic markers for Parkinson’s disease. Sci Rep. 2017;7:7328. doi: 10.1038/s41598-017-06767-y

27. Pell VR, Chouchani ET, Frezza C, Murphy MP, Krieg T. Succinate metabolism: a new therapeutic target for myocardial reperfusion injury. Cardiovasc Res. 2016;111:134–141. doi: 10.1093/cvr/cvw100

28. Jespersen NR, Hjortbak MV, Lassen TR, Stottrup NB, Johnsen J, Tonnesen PT, Larsen S, Kimose HH, Botker HE. Cardioprotective effect of succinate dehydrogenase inhibition in rat hearts and human myocardium with and without diabetes mellitus. Sci Rep. 2020;10:10344. doi: 10.1038/s41598-020-67247-4

29. Schulz R, Heusch G. Targeted Mito- and Cardioprotection by Malonate. Circ Res. 2022;131:542-544. doi: 10.1161/CIRCRESAHA.122.321582

30. Gallinat A, Mendieta G, Vilahur G, Padro T, Badimon L. DJ-1 administration exerts cardioprotection in a mouse model of acute myocardial infarction. Front Pharmacol. 2022;13:1002755. doi: 10.3389/fphar.2022.1002755

31. Billia F, Hauck L, Grothe D, Konecny F, Rao V, Kim RH, Mak TW. Parkinson-susceptibility gene DJ-1/PARK7 protects the murine heart from oxidative damage in vivo. Proc Natl Acad Sci U S A. 2013;110:6085–6090. doi: 10.1073/pnas.1303444110

32. Ong SB, Subrayan S, Lim SY, Yellon DM, Davidson SM, Hausenloy DJ. Inhibiting mitochondrial fission protects the heart against ischemia/reperfusion injury. Circulation. 2010;121:2012–2022. doi: 10.1161/CIRCULATIONAHA.109.906610

33. Maneechote C, Palee S, Kerdphoo S, Jaiwongkam T, Chattipakorn SC, Chattipakorn N. Differential temporal inhibition of mitochondrial fission by Mdivi-1 exerts effective cardioprotection in cardiac ischemia/reperfusion injury. Clin Sci (Lond). 2018;132:1669–1683. doi: 10.1042/CS20180510

34. Ding J, Zhang Z, Li S, Wang W, Du T, Fang Q, Wang Y, Wang DW. Mdivi-1 alleviates cardiac fibrosis post myocardial infarction at infarcted border zone, possibly via inhibition of Drp1-Activated mitochondrial fission and oxidative stress. Arch Biochem Biophys. 2022;718:109147. doi: 10.1016/j.abb.2022.109147

35. Rosdah AA, Abbott BM, Langendorf CG, Deng Y, Truong JQ, Waddell HMM, Ling NXY, Smiles WJ, Delbridge LMD, Liu GS, et al. A novel small molecule inhibitor of human Drp1. Sci Rep. 2022;12:21531. doi: 10.1038/s41598-022-25464-z

36. Bordt EA, Zhang N, Waddell J, Polster BM. The Non-Specific Drp1 Inhibitor Mdivi-1 Has Modest Biochemical Antioxidant Activity. Antioxidants (Basel). 2022;11. doi: 10.3390/antiox11030450

37. Kuznetsov AV, Javadov S, Sickinger S, Frotschnig S, Grimm M. H9c2 and HL-1 cells demonstrate distinct features of energy metabolism, mitochondrial function and sensitivity to hypoxia-reoxygenation. Biochim Biophys Acta. 2015;1853:276–284. doi: 10.1016/j.bbamcr.2014.11.015

38. Wang H, Song P, Du L, Tian W, Yue W, Liu M, Li D, Wang B, Zhu Y, Cao C, et al. Parkin ubiquitinates Drp1 for proteasome-dependent degradation: implication of dysregulated mitochondrial dynamics in Parkinson disease. J Biol Chem. 2011;286:11649–11658. doi: 10.1074/jbc.M110.144238

39. Li Y, Xiong Z, Jiang Y, Zhou H, Yi L, Hu Y, Zhai X, Liu J, Tian F, Chen Y. Klf4 deficiency exacerbates myocardial ischemia/reperfusion injury in mice via enhancing ROCK1/DRP1 pathway-dependent mitochondrial fission. J Mol Cell Cardiol. 2023;174:115–132. doi: 10.1016/j.yjmcc.2022.11.009

40. Bo T, Yamamori T, Suzuki M, Sakai Y, Yamamoto K, Inanami O. Calmodulin-dependent protein kinase II (CaMKII) mediates radiation-induced mitochondrial fission by regulating the phosphorylation of dynamin-related protein 1 (Drp1) at serine 616. Biochem Biophys Res Commun. 2018;495:1601–1607. doi: 10.1016/j.bbrc.2017.12.012

41. Du J, Li H, Song J, Wang T, Dong Y, Zhan A, Li Y, Liang G. AMPK Activation Alleviates Myocardial Ischemia-Reperfusion Injury by Regulating Drp1-Mediated Mitochondrial Dynamics. Front Pharmacol. 2022;13:862204. doi: 10.3389/fphar.2022.862204

42. Shou J, Huo Y. PINK1 Phosphorylates Drp1(S616) to Improve Mitochondrial Fission and Inhibit the Progression of Hypertension-Induced HFpEF. Int J Mol Sci. 2022;23. doi: 10.3390/ijms231911934

43. Hartmann S, Ridley AJ, Lutz S. The Function of Rho-Associated Kinases ROCK1 and ROCK2 in the Pathogenesis of Cardiovascular Disease. Front Pharmacol. 2015;6:276. doi: 10.3389/fphar.2015.00276

44. Brand CS, Tan VP, Brown JH, Miyamoto S. RhoA regulates Drp1 mediated mitochondrial fission through ROCK to protect cardiomyocytes. Cell Signal. 2018;50:48–57. doi: 10.1016/j.cellsig.2018.06.012

45. Arita R, Hata Y, Nakao S, Kita T, Miura M, Kawahara S, Zandi S, Almulki L, Tayyari F, Shimokawa H, et al. Rho kinase inhibition by fasudil ameliorates diabetes-induced microvascular damage. Diabetes. 2009;58:215–226. doi: 10.2337/db08-0762

46. Guo Y, Pu WT. Cardiomyocyte Maturation: New Phase in Development. Circ Res. 2020;126:1086–1106. doi: 10.1161/CIRCRESAHA.119.315862

47. Lu Q, Sun X, Yegambaram M, Ornatowski W, Wu X, Wang H, Garcia-Flores A, Da Silva V, Zemskov EA, Tang H, et al. Nitration-mediated activation of the small GTPase RhoA stimulates cellular glycolysis through enhanced mitochondrial fission. J Biol Chem. 2023;299:103067. doi: 10.1016/j.jbc.2023.103067

48. Hausenloy DJ, Yellon DM. New directions for protecting the heart against ischaemia-reperfusion injury: targeting the Reperfusion Injury Salvage Kinase (RISK)-pathway. Cardiovasc Res. 2004;61:448–460. doi: 10.1016/j.cardiores.2003.09.024

49. Karsdal MA, Nielsen SH, Leeming DJ, Langholm LL, Nielsen MJ, Manon-Jensen T, Siebuhr A, Gudmann NS, Ronnow S, Sand JM, et al. The good and the bad collagens of fibrosis - Their role in signaling and organ function. Adv Drug Deliv Rev. 2017;121:43–56. doi: 10.1016/j.addr.2017.07.014

50. Talman V, Ruskoaho H. Cardiac fibrosis in myocardial infarction-from repair and remodeling to regeneration. Cell Tissue Res. 2016;365:563–581. doi: 10.1007/s00441-016-2431-9

